# ppigFinder: an integrated desktop application for bacterial genome annotation and AlphaFold 3-based protein–protein interaction screening

**DOI:** 10.64898/2026.08.23.746524

**Authors:** Gabriel Umaji Oka, Camilla Adan, Celso Vítor Alves Queiroz Calomeno, Robson Francisco de Souza

## Abstract

**Motivation:** AlphaFold-based structure prediction has transformed structural biology by enabling accurate protein modelling and providing a powerful framework for inferring protein-protein interactions (PPIs). However, discovering candidate PPIs directly from genome sequences remains a fragmented and largely trial-and-error process, typically requiring separate tools for open reading frame (ORF) prediction, functional annotation, candidate selection, iterative testing of potential partners, manual preparation of individual structural-prediction jobs, and downstream interpretation of confidence metrics.

**Results:** We present Protein-Protein Interaction Genomic Finder (ppigFinder), a standalone, cross-platform desktop application that integrates these steps into a project-oriented graphical workflow for genome-based PPI discovery from nucleotide sequence data. ppigFinder combines ORF prediction, functional annotation, genomic-neighbourhood inspection, AlphaFold 3 job generation, remote job submission, and structural-confidence analysis within a single environment. As a proof of concept, we performed a VirD4-centered AlphaFold 3 interactome screen in Xanthomonas citri pv. citri strain 306, modelling VirD4 (ORF2601) against all 4,303 predicted chromosomal ORFs. Ranking by the minimum interchain predicted aligned error (PAE_min) placed all 14 XVIPCD-containing effector candidates within the top 1% of predictions, with the six top-ranked models corresponding to XVIP candidates. The screen also recovered an XVIPCD-containing protein absent from the reference genome annotation and identified high-confidence candidates predicted to bind VirD4 at a surface opposite to the XVIPCD-binding site.

**Availability and implementation:** ppigFinder is implemented in Python 3.11 and is freely available under the MIT licence at https://github.com/leepusp/ppigfinder, with documentation and installation instructions for Linux, macOS and Windows. The version described here is archived at [DOI Zenodo — XXXX].

## Introduction

Public databases contain a rapidly growing number of complete bacterial genome sequences, yet the protein interaction networks they encode remain largely uncharacterized. Starting from a newly assembled genome, the prediction of candidate PPIs still requires a series of loosely connected operations: ORF prediction, domain and homology-based annotation, candidate identification, preparation of structural-prediction inputs, one-pair-at-a-time submission of candidate complexes, and interpretation of the resulting models. These operations are typically performed across different applications, each with distinct input requirements, output formats and downstream processing constraints. Bridging ORF prediction, functional annotation, genomic context and structural PPI analysis within a single environment therefore remains an unmet need.

Genome-scale interactome prediction using AlphaFold-based approaches has already been demonstrated for several model organisms and biological systems. Reported applications include the essential interactomes of *Escherichia coli* and *Bacillus subtilis(Gómez Borrego and Torrent Burgas 2024)*, pooled AlphaFold 3 predictions for *Mycoplasma genitalium (Todor et al. 2025)*, proteome-scale screens of the human interactome (Schmid and Walter 2025), and pathogen interactome screens across 19 human bacterial pathogens using RoseTTAFold2-Lite (Humphreys et al. 2024). In parallel, dedicated tools have facilitated large-scale AlphaFold-based PPI investigations by automating structural prediction workflows, candidate screening and downstream confidence analysis (Yu et al. 2023; Rouger et al. 2025). These tools, however, are command-line oriented and take curated protein sets as input; none of them starts from a nucleotide sequence, and none integrates genomic context into candidate selection or interpretation.

Type IV secretion systems (T4SSs) are widespread and functionally diverse bacterial nanomachines that mediate conjugative DNA transfer, the delivery of virulence effectors into eukaryotic host cells, and the intoxication of competing bacteria (Alvarez-Martinez and Christie 2009; Sgro et al. 2019; Costa et al. 2024). T4SS-mediated secretion relies on VirD4 coupling proteins, which recognize cytoplasmic substrates and recruit them to the membrane-embedded translocation machinery (Llosa and Alkorta 2017). VirD4 belongs to the TrwB/TraG family of type IV coupling proteins, whose structural prototype, TrwB, assembles into a hexameric ring resembling ring helicases and F1-ATPase (Gomis-Rüth et al. 2001; 2002). Each protomer comprises an N-terminal membrane-associated region and a cytosolic nucleotide-binding ATPase domain containing an inserted all-alpha domain (AAD). In *Xanthomonas citri*, a yeast two-hybrid screen first identified a set of VirD4-interacting proteins (Alegria et al. 2005), which were subsequently recognized as antibacterial X-T4SS effectors sharing a conserved C-terminal *Xanthomonas* VirD4-interacting protein conserved domain (XVIPCD) (Souza et al. 2015). We later determined the solution NMR structure of the XVIPCD and demonstrated that it is directly recognized by the VirD4 AAD(Oka et al. 2022). This experimentally characterized system therefore provides a suitable benchmark for structure-assisted PPI discovery, with a defined positive set and a known binding mode.

Here we present Protein-Protein Interaction Genomic Finder (ppigFinder), a standalone, cross-platform desktop application that integrates established bioinformatic tools into a project-oriented graphical workflow for systematic genome-based PPI discovery directly from nucleotide sequence data **(Figure 1).** Within a visual genomic-context interface, ppigFinder combines modules for ORF prediction, conserved-domain and homology-based annotation, AlphaFold 3 input generation and structural-confidence analysis of predicted interactions. The AlphaFold 3 module supports several prediction-design strategies **(Figure 2):** hypothesis-driven, neighbourhood-guided, domain-guided, all-versus-all, genome-wide, and manually defined multi-chain analyses.

**Figure 1.**
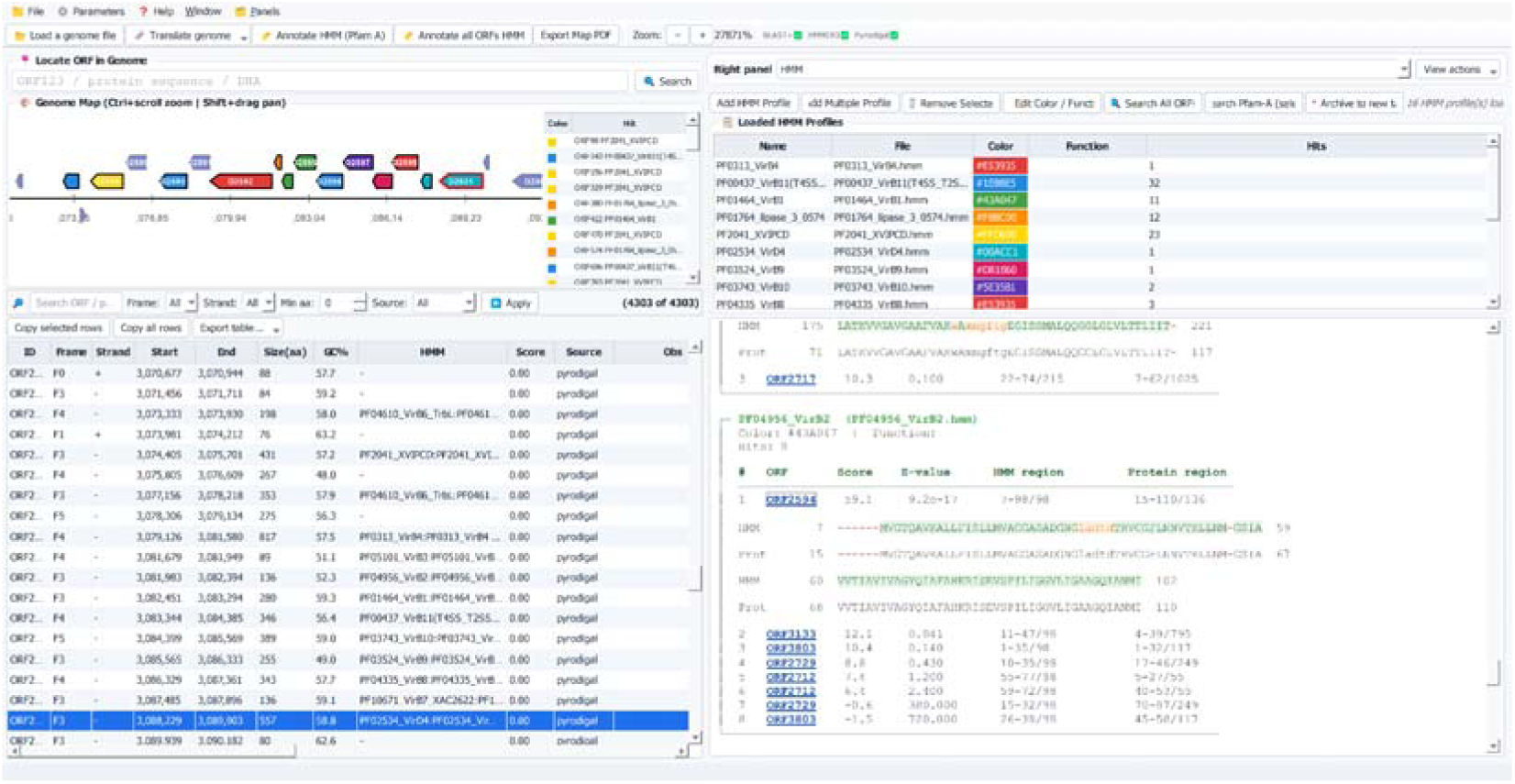
The ppigFinder graphical user interface for genome annotation and HMM profile management. The main window is organized into coordinated panels for genome visualization, ORF annotation and module-specific analysis. The top toolbar provides direct access to genome loading, ORF prediction, HMM-based annotation, whole-genome Pfam-A scanning (Mistry et al. 2021), genome-map export and zoom controls. In the example shown, the right-hand panel is set to the HMM profile analysis mode. The interactive genome map displays predicted ORFs as colour-coded directional arrows along a linear chromosomal coordinate axis, with colour corresponding to HMM profile hits defined in the loaded profile library. In the lower-left panel, the ORF annotation table summarizes the predicted ORFeome, listing ORF identifier, reading frame, strand, genomic coordinates, protein length, GC content, HMM annotation, HMM score, prediction source and user-defined observations. The selected ORF corresponds to a VirD4-like coupling protein annotated by HMMER3 profile search. The upper menu provides access to module-specific right-panel views: Domains (domain inspection), Neighborhood (genomic-context analysis), AlphaFold (prediction-input generation), Submit AF3 (remote submission and job monitoring), AF3 Table Results (ranking and inspection of prediction outputs), PPI Map (genome-wide visualization of predicted interaction pairs) and PPI Analysis (interface-centred analysis).

**Figure 2.**
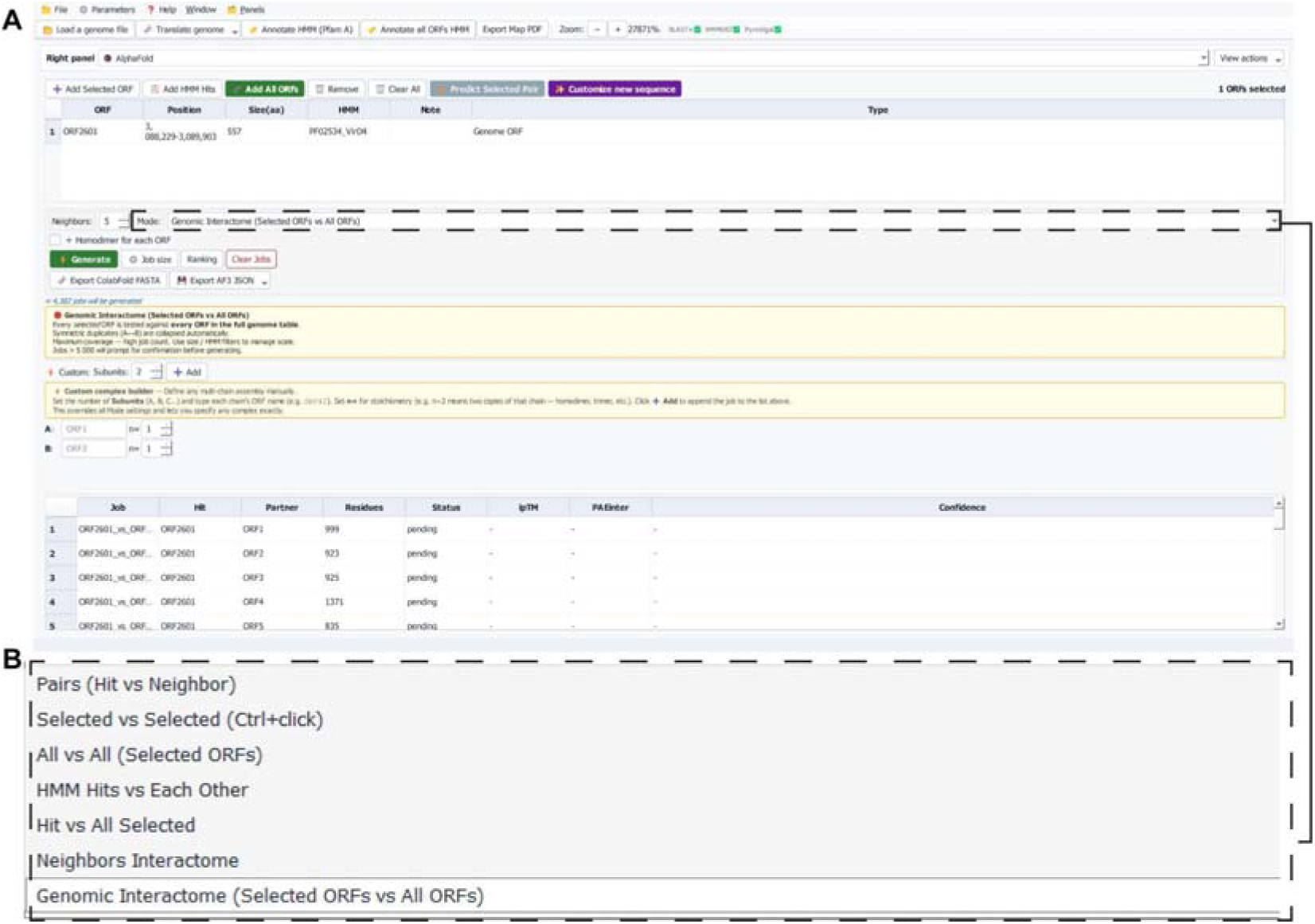
ppigFinder interface for AlphaFold 3 prediction-input generation. **(A)** AlphaFold 3 prediction-input panel. The ORF selection toolbar allows users to add individual ORFs, add HMM-annotated ORFs, add all predicted genome ORFs, remove entries, clear the selection, directly predict a selected pair, or add custom protein, RNA or DNA sequences. Custom sequence support enables the preparation of AlphaFold 3 inputs for protein–protein, nucleoprotein and protein–nucleic-acid complexes. The selection table lists each entry included in the prediction set, with ORF identifier, genomic position, protein size, HMM annotation, notes and chain type. In the example shown, the selected entry is ORF2601, corresponding to the VirD4 (KEGG: xac:XAC2623) coupling protein of the *Xanthomonas citri* pv. *citri* 306 antibacterial T4SS. The Neighbours parameter defines the number of flanking ORFs included in neighbourhood-based prediction modes. The Mode selector is set to Genomic Interactome (Selected ORFs vs All ORFs), in which each selected ORF is paired against all ORFs predicted in the genome; this mode was used to generate the VirD4-centred genome-wide prediction set, with symmetric duplicate pairs automatically collapsed. The estimated number of predictions is displayed before generation. The Generate button creates the prediction-input set, while additional controls allow token-limit configuration, ranking and export. Prediction inputs can be exported as AlphaFold 3-compatible JSON files or as ColabFold-compatible FASTA files (Mirdita et al. 2022). The Custom complex builder enables manual definition of multi-chain assemblies by specifying chain identifiers and stoichiometry, overriding the selected automated mode. **(B)** Interaction-mode selector. The Mode dropdown exposes multiple strategies for generating prediction inputs: *Pairs (Hit vs Neighbor)*, pairing selected ORFs with their nearest genomic neighbours; *Selected vs Selected*, testing manually selected ORF pairs; *All vs All*, generating all pairwise combinations within a selected set; *HMM Hits vs Each Other*, pairing ORFs identified by HMM profile searches; *Hit vs All Selected*, pairing HMM-hit entries against selected ORFs; *Neighbors Interactome*, generating a sliding-window neighbourhood interactome; and *Genomic Interactome*, pairing selected query ORFs against the complete predicted ORFeome. Together these modes enable hypothesis-driven, domain-guided, neighbourhood-guided, selected-set and genome-wide structural screening strategies from a single graphical interface.

We applied ppigFinder to the VirD4-centred interactome of the *X. citri* antibacterial T4SS, using VirD4 as the query in a genome-wide AlphaFold 3 screen against the 4,303 ORFs predicted from the X. citri 306 chromosome, thereby providing a computational counterpart to the original yeast two-hybrid screen. ppigFinder recovered all known XVIPCD-containing effectors within the top 1% of predictions, validating the screening strategy and supporting the prioritization of additional candidate VirD4 interactors that were not detected experimentally. Together, these analyses illustrate how ppigFinder integrates query-centered PPI screening, AlphaFold 3-derived interface-confidence metrics and genomic context to prioritize biologically relevant protein interactions directly from bacterial genome sequences.

## Methods

### Implementation

ppigFinder was implemented in Python 3.11. The graphical user interface was developed using PyQt6 (≥ 6.4), with automatic fallback to PyQt5 (≥ 5.15). Plotting of predicted aligned error (PAE) heatmaps and per-residue predicted local distance difference test (pLDDT) profiles, as well as figure export, was implemented using matplotlib ≥ 3.5 (Hunter 2007). Numerical and array operations were performed using NumPy ≥ 1.21 (Harris et al. 2020). Remote communication, file transfer and job submission to the DaVinci high-performance computing cluster were handled through SSH and SFTP using Paramiko ≥ 2.9 (Choi 2024). The application is distributed as source code and as PyInstaller-based executables for Linux, macOS and Windows.

### Genome data and ORF prediction

The complete chromosome sequence of *Xanthomonas citri pv. citri strain 306* (GenBank accession AE008923.1) was used throughout (da Silva et al. 2002). Open reading frames were predicted using Pyrodigal 2.x, a Python interface to the Prodigal gene-prediction algorithm (Hyatt et al. 2010; Larralde 2022), run in metagenomic mode with the bacterial and archaeal genetic code (translation table 11) and a minimum gene length of 30 codons (90 nt). Metagenomic mode is the ppigFinder default, which allows draft assemblies and single contigs to be processed without a separate training step, and was retained here for consistency with the standard workflow. Predicted ORFs were numbered sequentially along the chromosome (ORF0001-ORF4303) and translated in silico **(Supplementary Table 1).**

### Functional annotation

Protein-profile searches were performed in ppigFinder using the HMM annotation module, which runs HMMER3 ≥ 3.3 (Eddy 2011) to scan the translated ORFeome against a curated library of hidden Markov models (HMMs) of T4SS-associated protein families at a sequence E-value threshold of 0.01. All HMMs used in this study, including the XVIPCD profile (Pfam PF20410) used to identify XVIPCD-containing proteins, were downloaded from InterPro at the EMBL-EBI (Mistry et al. 2021). Sequence-similarity searches with BLASTp are also available in ppigFinder but were not used in the present analysis.

### AlphaFold 3 prediction-input generation

Prediction sets were generated using the AlphaFold 3 module of ppigFinder **(Figure 2).** For the genome-wide screen, the Genomic Interactome mode (Selected ORFs vs All ORFs) was used with VirD4 (ORF2601) as the single query, pairing it against every predicted ORF; symmetric duplicate pairs were automatically collapsed. Inputs were exported as AlphaFold 3-compatible JSON file **(Supplementary File 1)**, and can alternatively be exported as ColabFold-compatible FASTA files (Mirdita et al. 2022).

Three ORFs produced complexes exceeding the 3,000-token budget applied to the automated pipeline when combined with full-length VirD4 (557 residues): ORF1804/XAC1815 (4,743 residues), ORF2126/XAC2151 (3,397 residues) and ORF4150/XAC4213 (2,716 residues). These three complexes were instead submitted manually to the AlphaFold Server, which accepts up to 3000 tokens per job. ORF2126 and ORF4150 were modelled as full-length chains together with full-length VirD4, whereas ORF1804 exceeded the server limit as well and was split into two overlapping segments, each modelled with full-length VirD4: residues 1–3372 and residues 1372–4743, sharing an overlap of 2,001 residues. The final dataset therefore comprised 4,304 AlphaFold 3 models representing all 4,303 ORFs. ORF1804 was represented by two overlapping segment models instead of a single full-length model.

### Structure prediction

AlphaFold 3 predictions (Abramson et al. 2024) were generated on the DaVinci high-performance computing cluster at the Institute of Biomedical Sciences, University of São Paulo, using NVIDIA A30 and NVIDIA L40S GPU resources, providing approximately 24–48 GB of dedicated GPU memory (VRAM) per GPU, on compute nodes equipped with approximately either 1.0 or –1.5 TB of system RAM. Jobs were managed using the SLURM workload manager (Yoo et al. 2003). Unless otherwise specified, predictions were generated using the default AlphaFold 3 inference settings, with 10 recycles and five diffusion samples per random seed. The three complexes that exceeded the computational limits of the local installation, together with the VirD4 homodimer and homohexamer, were submitted manually to the AlphaFold Server (https://alphafoldserver.com) using the server default settings.

### Confidence metrics and candidate ranking

All predictions were analysed in ppigFinder, using the AF3 Table Results and PPI Analysis modules. ppigFinder scanned the AlphaFold 3 output directory of the whole ORFeome screen, parsing the summary_confidences.json and mmCIF files of each job folder, and automatically extracted PAE_min (Jumper et al. 2021; Abramson et al. 2024) and ipTM (Zhang and Skolnick 2004; Evans et al. 2021) for every predicted complex. PAE_min was taken as the minimum over both off-diagonal directions of the interchain PAE matrix. For graphical representation PAE_min was plotted as its reciprocal (1/PAE_min).

ppigFinder additionally computed a set of interface-centred descriptors for the 200 top-ranked predictions. Contact% is the percentage of token pairs in one off-diagonal block of the PAE matrix with PAE below 5 Å. The PAE hotspot delineates the confident contact patch: a rectangular window is anchored at the PAE minimum and then expanded or contracted edge by edge until it matches the extent of the low-error region, and is reported as its mean PAE, its size in residues, and a composite score combining the two. The best contact pair lists, for each chain, the residues closest to the partner chain in PAE terms, and the anchor windows give the amino-acid sequence flanking the PAE minimum in each chain.

## Results

The complete chromosome of *X. citri* pv. *citri* strain 306 was analysed using ppigFinder. ORF prediction with Pyrodigal identified 4,303 ORFs (Supplementary Table S1), which were subsequently screened against HMM profiles associated with T4SS components. The profile-based analysis recovered the known components of the chromosomal X-T4SS gene cluster, including the VirD4 coupling protein, assigned to ORF2601 and corresponding to the annotated locus XAC2623 (KEGG: xac:XAC2623). The only component not recovered under the default Pyrodigal metagenomic mode was XAC2611, a small X-T4SS-associated lipoprotein encoded within the *vir* locus locus (Oka et al. 2026; Alegria et al. 2005). Screening against the XVIPCD HMM profile identified 14 proteins containing the domain **(Table 1).** Twelve had been reported by Alegria *et al*. (2005) and a thirteenth by Souza *et al*. (2015). The fourteenth, ORF0763, is described here for the first time: a previously unreported XVIPCD-containing protein, absent from the reference KEGG annotation of the *X. citri* 306 genome (da Silva et al. 2002).

**Table 1.**
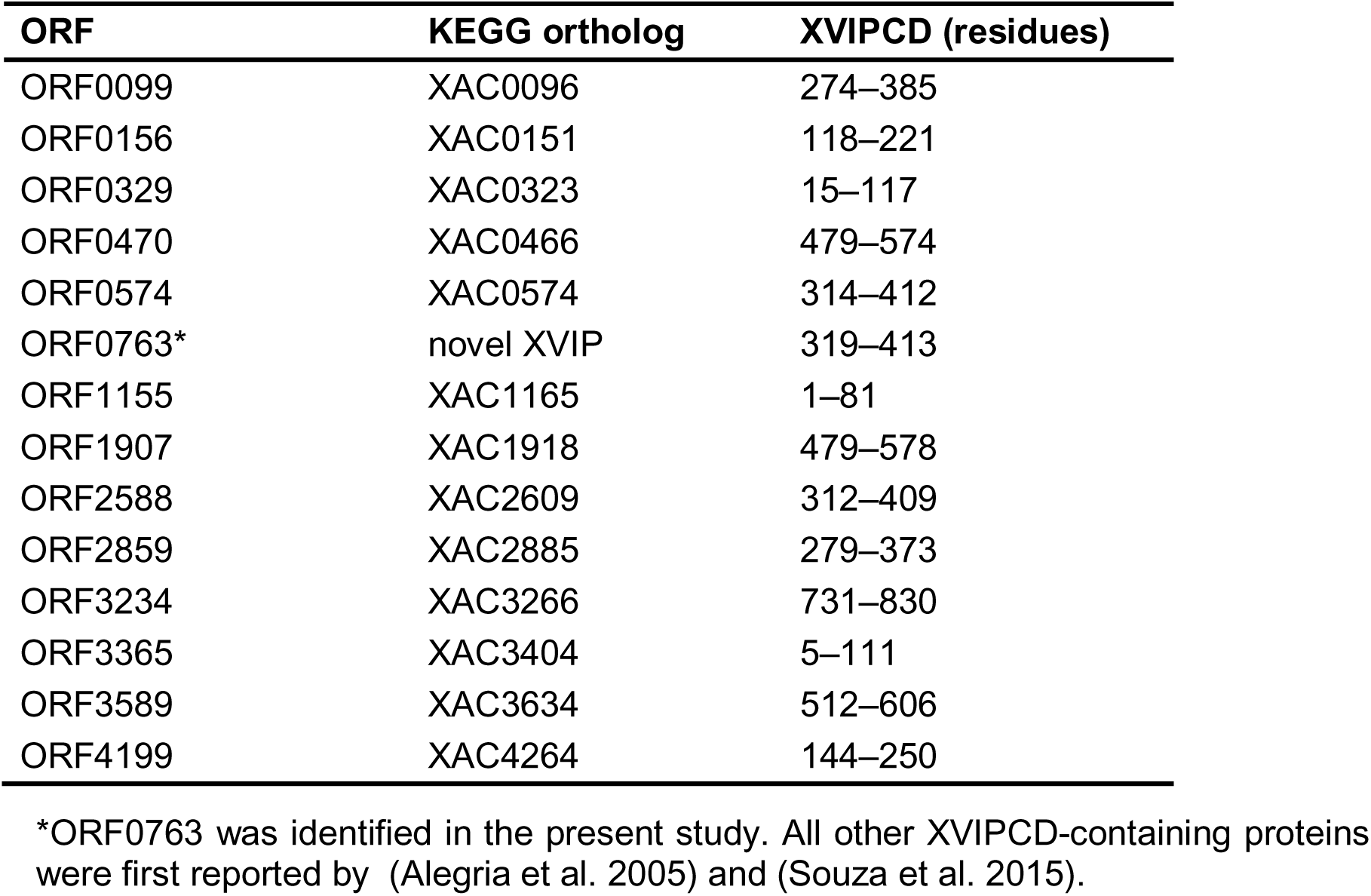
XVIPCD-containing proteins encoded in the *Xanthomonas citri* pv. *citri* strain 306 chromosome. ORFs were predicted with ppigFinder and screened against the XVIPCD HMM profile (Pfam PF20410). XVIPCD coordinates indicate amino acid positions within each predicted protein.

| ORF | KEGG ortholog | XVIPCD (residues) |
| --- | --- | --- |
| ORF0099 | XAC0096 | 274–385 |
| ORF0156 | XAC0151 | 118–221 |
| ORF0329 | XAC0323 | 15–117 |
| ORF0470 | XAC0466 | 479–574 |
| ORF0574 | XAC0574 | 314–412 |
| ORF0763* | novel XVIP | 319–413 |
| ORF1155 | XAC1165 | 1–81 |
| ORF1907 | XAC1918 | 479–578 |
| ORF2588 | XAC2609 | 312–409 |
| ORF2859 | XAC2885 | 279–373 |
| ORF3234 | XAC3266 | 731–830 |
| ORF3365 | XAC3404 | 5–111 |
| ORF3589 | XAC3634 | 512–606 |
| ORF4199 | XAC4264 | 144–250 |

\*ORF0763 was identified in the present study. All other XVIPCD-containing proteins were first reported by (Alegria et al. 2005) and (Souza et al. 2015).

### The genome-wide screen recovers the complete set of known VirD4 partners

The resulting VirD4–ORF predictions were ranked in ppigFinder (**Supplementary Table 2)**. The resulting VirD4–ORF predictions were ranked using ppigFinder (Supplementary Table S2). The genome-scale screen, comprising 4,304 models representing 4,303 ORFs, required approximately 1,076 aggregate GPU-hours (corresponding to 15 GPU-minutes per model or 250 GPU-hours per 1,000 models), which was efficiently distributed across parallel GPU nodes to complete the screen in approximately 11 days of wall-clock time, excluding queue time and CPU-based data preparation. Ranking by PAE_min revealed a pronounced enrichment of XVIPCD-containing proteins among the highest-confidence predictions **(Figure 3A)**. The six top-ranked predictions all corresponded to XVIP candidates, with PAE_min values between 1.16 and 1.38 Å, and XVIPs accounted for 11 of the top 20. All 14 XVIP candidates fell within the top 1% of the 4,304 models, with ipTM values between 0.60 and 0.82. The lowest-ranking of them, ORF2859/XAC2885, ranked 42nd overall (PAE_min 2.99 Å, ipTM 0.60).

**Figure 3.**
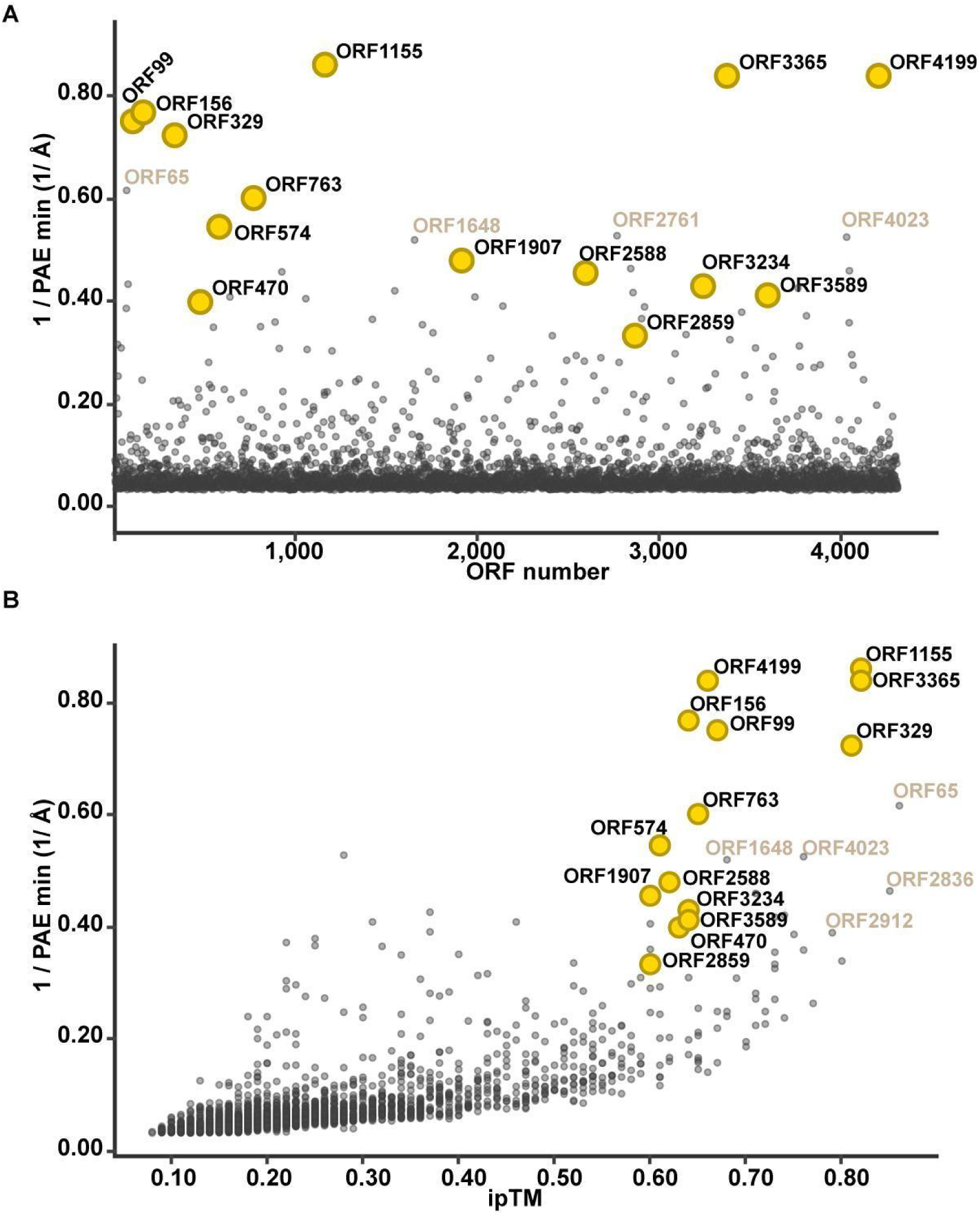
Genome-wide AlphaFold 3 screening identifies XVIPCD-containing proteins among the highest-ranking predicted VirD4 partners. **(A)** Scatter plot of the inverse minimum interchain predicted aligned error (1/PAE_min) for AlphaFold 3 models of VirD4 (ORF2601) in complex with each chromosomal ORF predicted in *Xanthomonas citri* pv. *citri* strain 306. The *x*-axis indicates the sequential ORF number along the chromosome. The 14 XVIPCD-containing candidates identified by HMM profile analysis are highlighted in yellow and labelled. Selected non-XVIP proteins with comparatively high 1/PAE_min values are labelled in brown. **(B)** Scatter plot of 1/PAE_min against ipTM for the same set of predicted VirD4–ORF complexes. XVIPCD-containing proteins are shown in yellow, selected high-scoring non-XVIP proteins in brown, and all remaining predictions in grey. The plots were generated via the PPI Analysis module of ppigFinder.

Several of the highest-ranking proteins consisted predominantly of an XVIPCD and lacked a recognizable N-terminal toxin domain, including ORF1155 (XAC1165; XVIPCD at residues 1–81), ORF3365 (XAC3404; residues 5–111) and ORF0329 (XAC0323; residues 15–117), further supporting the XVIPCD as the principal VirD4-recognition module. The compact architecture of these short proteins, which lack the large accessory domains present in most other candidates, may have contributed to the confidence of their predictions. This does not account for the observed enrichment, however, since XVIP candidates of up to 861 residues (ORF3234) also ranked within the top 20.

### Combining PAE_min with ipTM separates candidates from the background

Another metric we employed was ipTM, the interface predicted TM-score reported by AlphaFold 3 (Evans et al. 2021), which assesses the overall relative arrangement of the two chains rather than the confidence of a single contact. Because the two measures report complementary information, plotting 1/PAE_min against ipTM concentrated the XVIP candidates in the upper-right region of the distribution **(Figure 3B)**, a representation that proved useful for visual triage of the full screen.

### Conserved XVIPCD–AAD binding mode

Structural inspection of the predicted complexes indicated that a β-sheet within the XVIPCD of all 14 candidates engages the all-alpha domain (AAD) of VirD4 **(Figure 4)**, consistent with the binding mode previously characterized by solution NMR, isothermal titration calorimetry, pull-down assays and *in vivo* bacterial competition assays (Oka et al. 2022). Superposition of the 14 models shows a common orientation of the XVIPCD relative to the AAD **(Figure 4A)**, and the corresponding PAE maps display localized low-error patches confined to the off-diagonal quadrants at the XVIPCD position **(Figure 4D, E)**, the signature of a focal, domain-limited interface.

**Figure 4.**
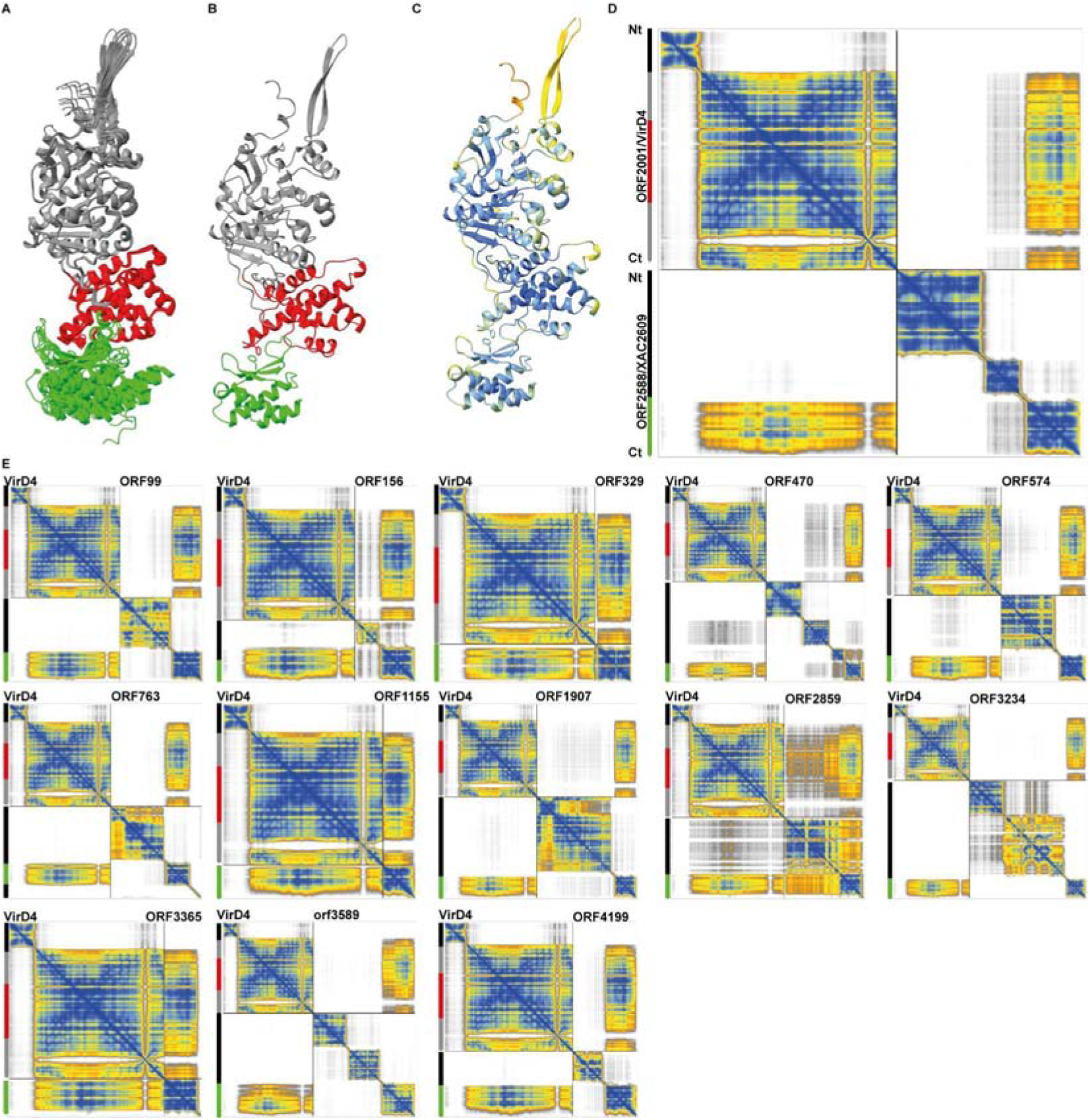
AlphaFold 3 models of VirD4 in complex with XVIPCD-containing proteins. **(A)** Superposition of the 14 AlphaFold 3 models of VirD4 in complex with XVIPCD-containing proteins. The VirD4 nucleotide-binding domain is shown in grey and the all-alpha domain (AAD) in red; the XVIPCD region of each partner is shown in green. The N-terminal transmembrane helices of VirD4, the predicted toxin domains and the final 20 C-terminal residues of each partner were omitted for clarity, to facilitate comparison of the conserved VirD4–XVIPCD binding mode. **(B)** AlphaFold 3 model of the predicted VirD4–ORF2588/XAC2609 complex. **(C)** The same complex coloured by residue-level AlphaFold 3 confidence, with blue indicating higher-confidence regions and yellow to orange indicating lower-confidence regions. **(D)** Predicted aligned error (PAE) map for the VirD4–ORF2588/XAC2609 complex. Schematic bars to the left and above the map indicate the N-to-C-terminal organization of each chain, with the N termini positioned at the top and left, respectively. For the XVIP partner, the XVIPCD is shown in green and the remaining regions in black. The diagonal quadrants report confidence in the relative positioning of residues within each chain, whereas the off-diagonal quadrants report confidence in the relative positioning of residues between the two chains. **(E)** PAE maps for VirD4 in complex with the remaining XVIPCD-containing proteins: ORF0099, ORF0156, ORF0329, ORF0470, ORF0574, ORF0763, ORF1155, ORF1907, ORF2859, ORF3234, ORF3365, ORF3589 and ORF4199.

### High-confidence candidates lacking a detectable XVIPCD

Several proteins not classified as XVIPCD-containing effectors combined low PAE_min values with relatively high ipTM scores, including ORF0065, ORF1539, ORF1648, ORF2761, ORF2836, ORF4023 and ORF4038 **(Figures 3 and 5; Supplementary Table 2)**. ORF0065 ranked 7th overall (PAE_min 1.62 Å, ipTM 0.86) and ORF2836 14th (PAE_min 2.15 Å, ipTM 0.85), the two highest ipTM values in the top 20. Structural inspection revealed structurally heterogeneous binding modes among these candidates **(Figure 5)**. In particular, ORF0065 and ORF4023 were predicted to bind closely to the VirD4 AAD at a surface opposite the XVIPCD-binding site.

**Figure 5.**
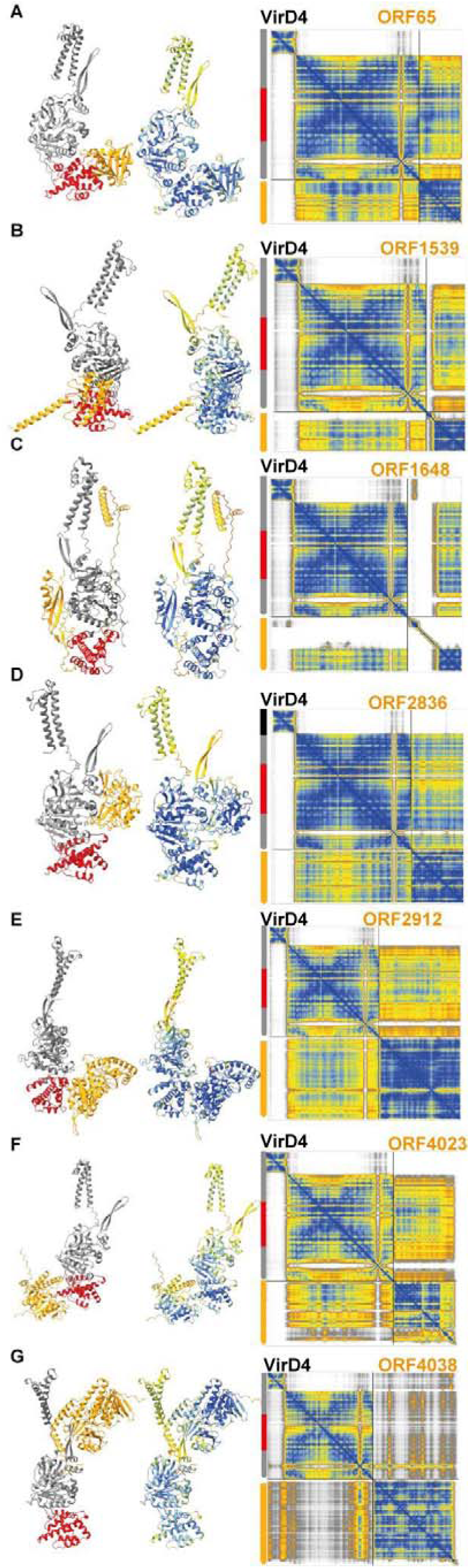
Structural inspection of selected high-scoring VirD4 partners lacking a detectable XVIPCD. (A–G) (A–G) AlphaFold 3 predictions of VirD4 in complex with ORF0065, ORF1539, ORF1648, ORF2761, ORF2836, ORF4023 and ORF4038, respectively. For each candidate, the model on the left is coloured by protein and VirD4 domain: the VirD4 nucleotide-binding domain (NBD) is shown in grey, the all-alpha domain (AAD) in red and the candidate partner in orange. The central model shows the same complex coloured according to residue-level AlphaFold 3 confidence (pLDDT), with blue indicating higher confidence and yellow to orange indicating lower confidence. The corresponding predicted aligned error (PAE) map is shown on the right. Schematic bars adjacent to each map illustrate the N- to C-terminal organization of VirD4 (top to bottom), highlighting the N-terminal transmembrane region and nucleotide-binding domain (NBD) in grey, and the all-alpha domain (AAD) in red, alongside the candidate protein shown in orange.

## Discussion

As a proof of concept, ppigFinder identified XVIPCD-containing proteins among the strongest predicted VirD4-interacting partners encoded in the *X. citri* genome, consistent with the VirD4–XVIP interactions previously identified experimentally by (Alegria et al. 2005). Recovery of this known positive set provides an important initial validation of the screening strategy. Notably, the workflow also identified ORF0763, a previously unannotated XVIPCD-containing candidate. The chromosome of *X. citri* pv. *citri* strain 306 (da Silva et al. 2002) contains 4,314 annotated protein-coding genes in the current KEGG reference annotation (organism *xac)*, a number comparable to the 4,303 ORFs predicted here; nevertheless, the two protein sets do not completely overlap. Although the yeast two-hybrid screen performed by (Alegria et al. 2005) used a prey library derived from total genomic DNA and could therefore, in principle, have sampled this genomic region independently of protein annotation, ORF0763 was not recovered for reasons that remain unclear. By contrast, because the PSI-BLAST analysis performed by (Souza et al. 2015) searched for annotated protein sequences, ORF0763 was not represented in the sequence space examined in that study.

ppigFinder was developed to make structure-based PPI screening accessible to users with a biological background but limited computational experience, including undergraduate students, while allowing the scale of the analysis to be tailored to the biological question and available hardware. Although exhaustive interactome screens require substantial computational infrastructure and may demand advanced computational expertise, focused analyses targeting a specific protein or protein family can be performed using consumer-grade or previous-generation GPUs (Yu et al. 2023; Gómez Borrego and Torrent Burgas 2024; Rouger et al. 2025).

AlphaFold 3 reports the minimum chain-pair PAE as a confidence measure that correlates with whether two chains interact and may, in some cases, distinguish binders from non-binders. Following this guidance, we employed this value, hereafter referred to as PAE_min, as the primary metric for ranking the VirD4–ORF predictions (Abramson et al. 2024). When predictions were ordered by PAE_min, the six highest-ranked models were XVIPs, 11 of the 14 XVIPs occurred among the top 20, and all 14 were recovered within the first 42 predictions. By comparison, ranking by either ipTM or the native AlphaFold 3 ranking score placed only three XVIPs among the top 20, with the complete set recovered within the first 71 and 77 predictions, respectively **(Supplementary Table 2).** The mean interchain PAE was less discriminative: within the subset of 208 models for which full PAE matrices were available, its median was similar for XVIP and non-XVIP predictions (25.51 and 24.77 Å, respectively).

The models themselves can be checked against what is already known about this interface. Structural inspection indicated that a β-sheet within the XVIPCD of all 14 candidates engages the AAD of VirD4 **(Figure 4)**, and the anchor windows computed by ppigFinder locate that contact in sequence. On the VirD4 side, all 14 candidates anchored within residues 238 to 304, entirely inside the AAD (residues 197 to 355) that was shown by yeast two-hybrid, isothermal titration calorimetry and NMR to mediate XVIPCD recognition (Oka et al. 2022). On the effector side, the correspondence is finer. ORF2588 corresponds to X-Tfe^XAC2609^, the protein whose XVIPCD structure was solved, and its predicted anchor window spans residues 374 to 394, containing Phe375, Val377, Asp383 and Pro384. These are the residues whose amide resonances shift in addition of VirD4_AAD_ and whose substitution reduces or abolishes binding, the F375A/V377A double mutant showing no detectable interaction by calorimetry. A window derived solely from predicted aligned error therefore converges on the residues identified experimentally, which supports using anchor windows to nominate positions for mutagenesis in candidates that have not yet been characterized. The models nevertheless remain theoretical, and experimental structures will be required to confirm the predicted interfaces and define the molecular basis of substrate coupling.

Conversely, X-T4SS components including VirB4, VirB10 and VirB11 yielded low-confidence predicted interfaces against VirD4, despite experimental evidence for their association with VirD4-like coupling proteins in other T4SSs (Llosa et al. 2003; Atmakuri et al. 2004; Ripoll-Rozada et al. 2013). Manual inspection of the VirD4–VirB10 model revealed that the N-terminal transmembrane helix of VirD4 was positioned close to an N-terminal transmembrane helix of **VirB10 (Supplementary Figure 1)**. This illustrates how automated ranking based solely on confidence metrics over pairwise interactions may deprioritize localized, stoichiometry- or assembly-dependent interfaces. VirD4 itself provides a clearer demonstration of this limitation: the VirD4 dimer was predicted poorly, whereas specifying six copies produced a confident hexameric ring consistent with the TrwB structure **(Supplementary Table 3)** (Gomis-Rüth et al. 2001).

Beyond the canonical XVIPCD-containing proteins, the screen identified several promising VirD4 partner candidates that were not recovered in the earlier yeast two-hybrid study **(Figure 5)**. When full-length VirD4 was used as bait, all 68 sequenced preys were derived from 12 XVIP-containing proteins and contained the XVIPCD (Alegria et al. 2005). The genomic prey fragments were expressed downstream of the Gal4 activation domain, generating Gal4AD–prey fusions in which the prey sequences occupied the C-terminal position (Alegria et al. 2005; Uetz and Hughes 2000). This configuration placed C-terminal interaction regions distal from the fusion junction and may have favored the recovery of XVIPCD-mediated interactions, as fusion orientation can substantially influence the interactions detected in yeast two-hybrid assays (Stellberger et al. 2010). By contrast, the AlphaFold 3 screen imposed no fusion junction or predefined interaction region and therefore allowed alternative binding modes to be explored. Notably, ORF0065/XAC0064 and ORF4023/XAC4088 showed low interchain PAE_min values and ipTM scores comparable to, or more favorable than, those of several XVIPCD-containing candidates. Both proteins were predicted to contact the VirD4 AAD at a surface opposite the established XVIPCD-binding site, revealing structurally distinct candidate interfaces for experimental validation.

## Conclusion

We benchmarked ppigFinder’s AF3-based approach in a genome-wide screen starting from unannotated genomic sequence, using a T4SS effector family with an experimentally defined positive set as a reference. The graphical interface provides access to complex embedded pipelines and to ranking metrics essential for rapidly generating and evaluating several thousand pairwise interaction models, and supports workflows not exercised here, including individual pair testing, multi-chain assemblies of defined stoichiometry, manual curation of ORF annotations, and inspection of sequence features and genomic context alongside the structural predictions. ppigFinder is therefore suited to poorly characterized or recently isolated genomes, and to groups that use structural predictions to generate testable hypotheses for experimental validation. Future developments will enable other protein complex prediction methods and the integration of alignment-based covariation filters, which should speed up the analysis while preserving its sensitivity.

## Acknowledgements

The authors thanks the CEPID B3. The performance computing resources were provided by CEPID B3, and additional tests were performed using the DaVinci server at ICB-USP.

## Data availability

The *Xanthomonas citri* pv. *citri* strain 306 chromosome sequence is available from GenBank under accession AE008923.1. All AlphaFold 3 confidence metrics generated in this study are provided in Supplementary Tables 2 and 3. Structural models can be provided by request.

## Code availability

ppigFinder is freely available at https://github.com/leepbioinfo/ppigfinder under the MIT licence. The version described here is archived at **[DOI Zenodo —XXX]**.

## Supplementary Tables — legends and column descriptions

**Supplementary Table 1.**
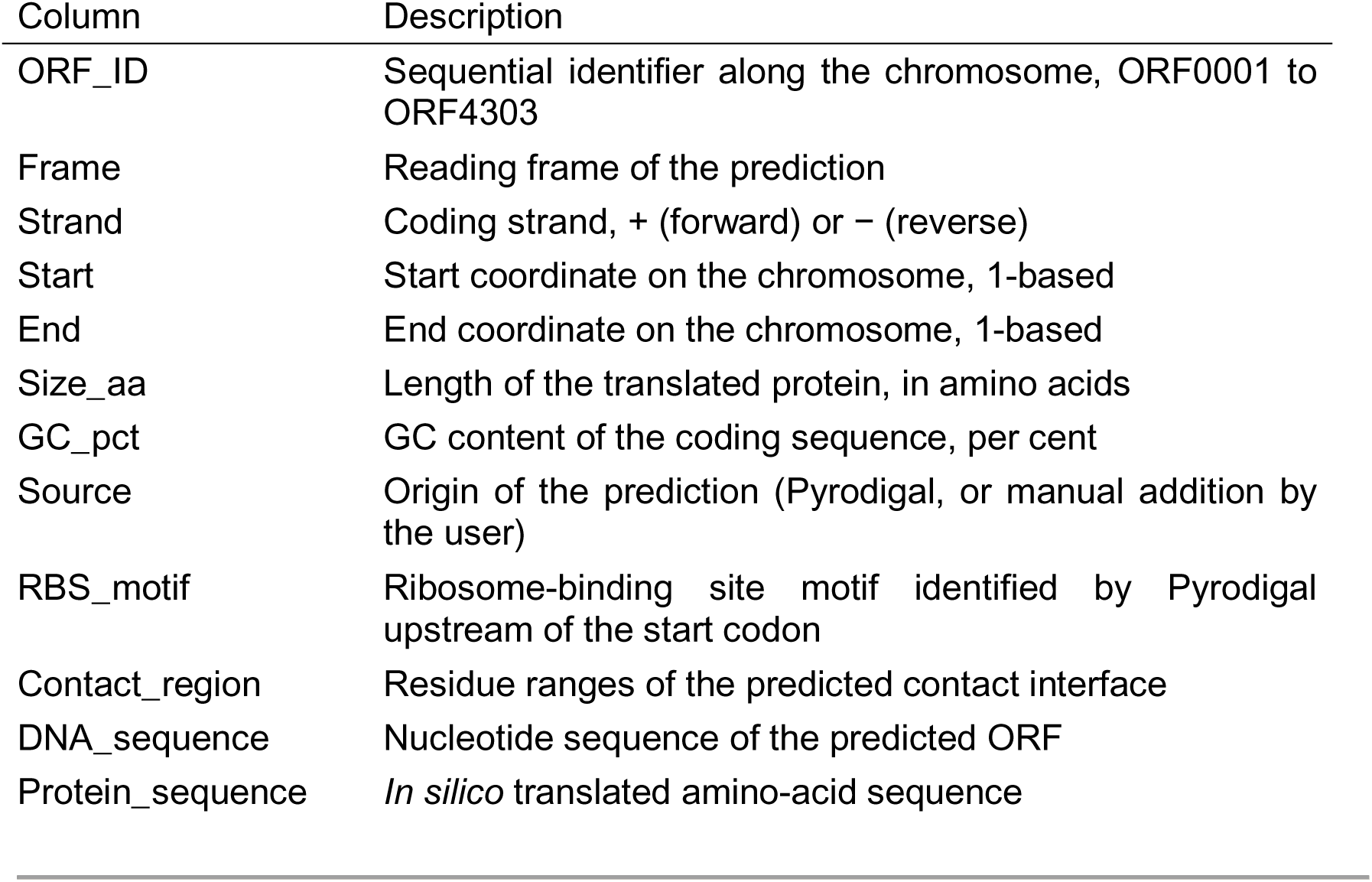
ppigFinder predicted ORFeome of *Xanthomonas citri* pv. *citri* strain 306 Complete set of open reading frames predicted from the *X. citri* pv. *citri* 306 chromosome (GenBank AE008923.1) with Pyrodigal, together with the functional annotation assigned by ppigFinder and the *in silico* translation of each ORF. ORFs are numbered sequentially along the chromosome (ORF0001–ORF4303). The table is the export of the central ORF panel of ppigFinder and contains one row per predicted ORF.

| Column | Description |
| --- | --- |
| ORF_ID | Sequential identifier along the chromosome, ORF0001 to ORF4303 |
| Frame | Reading frame of the prediction |
| Strand | Coding strand, + (forward) or – (reverse) |
| Start | Start coordinate on the chromosome, 1-based |
| End | End coordinate on the chromosome, 1-based |
| Size_aa | Length of the translated protein, in amino acids |
| GC_pct | GC content of the coding sequence, per cent |
| Source | Origin of the prediction (Pyrodigal, or manual addition by the user) |
| RBS_motif | Ribosome-binding site motif identified by Pyrodigal upstream of the start codon |
| Contact_region | Residue ranges of the predicted contact interface |
| DNA_sequence | Nucleotide sequence of the predicted ORF |
| Protein_sequence | *In silico* translated amino-acid sequence |

**Supplementary Table 2.**
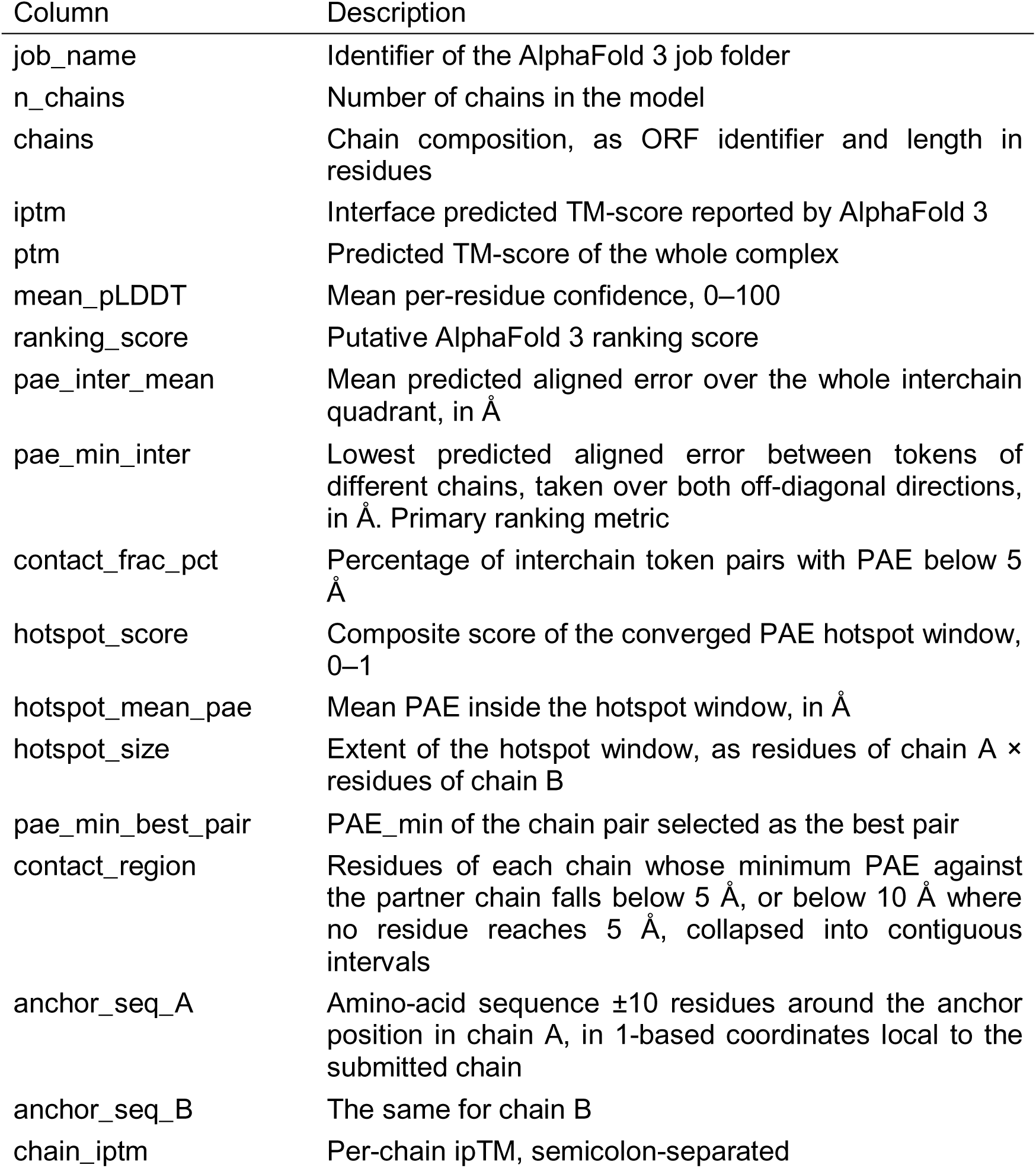
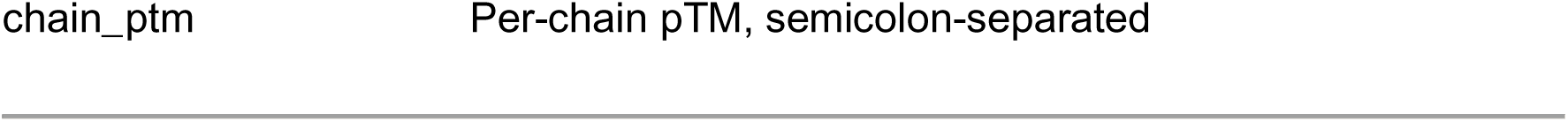
Supplementary Table S2. ppigFinder AlphaFold 3 confidence metrics for the VirD4-centered interactome scree. Confidence metrics for every model generated in the genome-wide screen, in which VirD4 (ORF2601 / XAC2623) was paired with each ORF of the predicted ORFeome. One row per model. Global metrics (iptm, ptm, ranking_score, pae_min_inter, contact_frac_pct) were extracted for all models. Interface-centred descriptors (mean_pLDDT, pae_inter_mean, the three hotspot_* fields, contact_region and the two anchor sequences) require parsing of the full PAE matrix and were computed for the 200 top-ranked models. Empty cells indicate metrics that were not computed for that model rather than absent values.

**Supplementary Table 3.** AlphaFold 3 confidence metrics for VirD4 self-association VirD4 (ORF2601 / XAC2623) modelled as a homodimer and as a homohexamer. Both models were generated on the AlphaFold Server, and all values are read directly from the summary_confidences.json file of each job. The homodimer was also generated independently as part of the genome-wide screen on the DaVinci cluster (Supplementary Table S2), which gave ipTM 0.14 and PAE_min 26.31 Å, in agreement with the values reported here.

| Row | Description |
| --- | --- |
| Chains modelled | Number of full-length VirD4 chains (557 residues each) in the assembly |
| Total residues (tokens) | 557 × number of chains |
| Unique interchain pairs | $n(n-1)/2$, the number of distinct chain-pair interfaces |
| ipTM | Interface predicted TM-score of the assembly |
| ptm | Predicted TM-score of the whole assembly |
| ranking_score | Native AlphaFold 3 ranking score |
| chain_ipTM | ipTM computed for each individual chain; identical across all chains in both models |
| chain_ptm | pTM computed for each individual chain; identical across all chains in both models |
| chain_pair_ipTM, lowest and highest | Range of the off-diagonal entries of the chain_pair_ipTM array, across all chain-pair interfaces |
| PAE_min, lowest, median and highest | Distribution of the minimum interchain predicted aligned error across all chain pairs, taken over both off-diagonal directions, in Å |
| fraction_disordered | Fraction of the assembly predicted to be disordered |
| has_clash | 0 indicates that no steric clash was detected |

